# Generation of an induced pluripotent stem cell line ERCi003-A derived from a patient with maturity-onset diabetes of the young type 10 caused by a heterozygous *INS* mutation

**DOI:** 10.1101/2024.12.04.626760

**Authors:** Asya Bastrich, Daniil Antonov, Aleksandra Podzhilkova, Darya A. Petrova, Svetlana V. Pylina, Dmitriy N. Laptev, Elena A. Sechko, Sergey N. Kuznetsov, Ekaterina A. Vetchinkina, Natalia G. Mokrysheva

## Abstract

Maturity-onset diabetes of the young type 10 (MODY10) is inherited in an autosomal dominant manner, characterized by islet cell dysfunction, impaired insulin synthesis and secretion. MODY10 is relatively rare, and is caused by a mutation of the 11p15.5 site on chromosome 11 encoding insulin. We obtained iPSCs (ERCi003-A line) from dermal fibroblasts derived from a patient with MODY10 carrying a previously undescribed pathogenic heterozygous mutation in the *INS* gene (c.93C>G, p.C31W) into iPSCs using transfection with self-replicating RNA vector. iPSCs proliferate in dense monolayer cell colonies, have a normal karyotype (46,XY), express pluripotency markers (OCT4, SOX2, TRA-1-60). The functional pluripotency of iPSCs was confirmed by their ability to form embryoid bodies and differentiate into the three germ layers (ecto-, endo-, and mesoderm). Sanger sequencing of iPSCs confirmed the presence of pathogenic heterozygous mutation in the *INS* gene. This cell line could be useful to thoroughly modeling of MODY10 to improve clinical understanding of the disease.

## INTRODUCTION

MODY is a both clinically and genetically heterogeneous group of monogenic disorders characterized by β-cell dysfunction, autosomal dominant inheritance, onset at a young age (mainly in the second or third decades), absence of autoantibodies specific for type 1 diabetes mellitus and by absence of signs of metabolic syndrome. MODY-causing mutations have been identified in 14 different genes (Greeley et al., 2022). As it is known preproinsulin is synthesized by the transcription and translation of *INS*, and subsequently cleaved to secrete insulin. Hence, *INS* mutations are strongly associated with abnormal insulin generation and glucose metabolism. In most cases mutations of the 11p15.5 site on chromosome 11 encoding insulin are the cause of permanent neonatal diabetes mellitus. However, it has now become known that a mutation in this gene can also lead to the development of MODY10 (Edghill et al., 2008; Molven et al., 2008).

Here, we reported the iPSC line from a patient carrying a heterozygous mutation (c.93C>G, p.C31W) in the *INS* gene which is associated with MODY10. This mutation was recently described for the first time by the clinical specialists of our center as a part of a family medical history (Sechko et al., 2022). MODY10 is uncommon, especially when caused by the c.93C>G mutation, and thus, our knowledge of this disease is limited. There are only a few descriptions of families with MODY-INS or MODY10. Patient-specific iPSC line carrying mutations in the *INS* gene c.93C>G can be used for study of molecular mechanisms of MODY10 pathology in the isogenic system by genome editing and differentiation into stem-cell-derived pancreatic islets (Balboa et al., 2022). Edited iPSCs can serve as a basis for the establishment of personalized therapies for INS-associated MODY10 and improving current therapeutic approaches.

## MATERIALS AND METHODS

### Titration of the antibiotic

Human dermal fibroblasts (HDFs) were seeded at a density of 3,000 cells/cm^2^ in a 24-well plate in DMEM (Gibco, USA) containing 10% FBS (Cytiva, USA), 1x GlutaMAX (Gibco), 1x NEAA (Gibco). Upon reaching 50–60% confluence, the cells were treated with puromycin dihydrochloride (Thermo Fisher Scientific, USA) for 5 days with daily media changes to maintain the antibiotic concentration. The antibiotic concentration range was from 0.2 µg/ml to 0.48 µg/ml with increasing 0.02 µg/ml interval. Cell viability was evaluated after five full days of selection. The concentration of the antibiotic resulting in 80% cell death (IC80) was determined to be 0.36 µg/ml.

### iPSCs reprogramming and cell culture

HDFs were transfected using self-replicating RNA vector ReproRNA™-OKSGM (Stem Cell Technologies, Canada) according to the manufacturer’s instruction followed by puromycin (Thermo Fisher Scientific) selection. On the 21st day of post-transfection, colonies were picked and cells were seeded in a 96-well plate coated with Matrigel (Corning, USA). Long-term feeder free culture of ERCi003-A iPSCs was conducted in mTeSR™1 (Stem Cell Technologies) under 37°C, 5% CO2. iPSCs were passaged with Versene solution (Paneco, Russia) at a ratio of 1:15 – 1:20 every 4–5 days with ROCK-inhibitor (Stem Cell Technologies).

### Immunocytochemistry

The immunocytochemistry method was used to assess the pluripotency of ERCi003-A at the 14th passage. Cells were fixed with 4% paraformaldehyde (Thermo Fisher Scientific) at room temperature (RT) for 15 minutes, washed twice with DPBS (Gibco), permeabilized with 0.5% Triton-X100 for 10 mins and then blocked with PBST containing 5% FBS (Cytiva) for 2 hours. Primary antibodies were incubated, at the concentrations listed in Table 1, in 2.5% FBS solution overnight at 4ºC. Next day, the cells were washed with DPBS thrice and then exposed to corresponding secondary antibodies for 1 hour at RT. For nuclei staining DAPI (Thermo Fisher Scientific) in concentration 1 µg/ml was used. The images were obtained and analyzed by fluorescence microscopy using Axio Observer 7 equipped with a Colibri 7 (Zeiss, Germany).

**Table 1.**
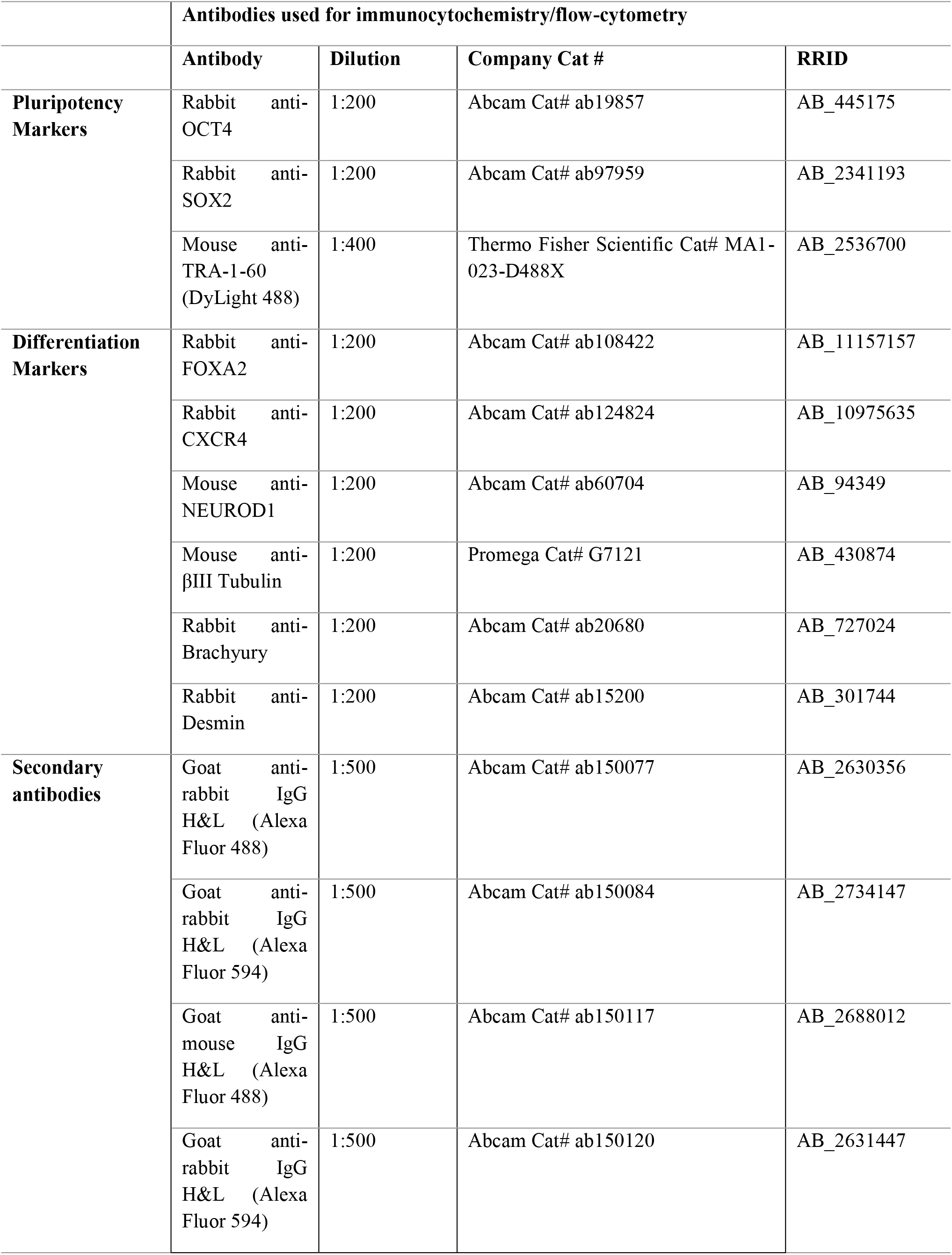
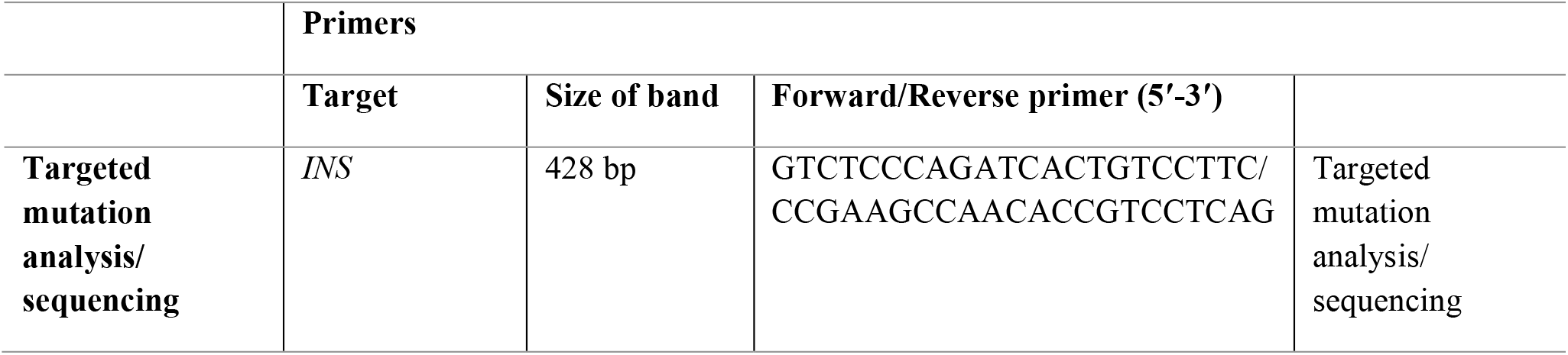
Reagents details.

### Flow cytometry

iPSC colonies were collected at a high confluence of up to 80% at the 13th passage. Cells were dissociated into single cell suspension using TrypLE Express (Thermo Fisher Scientific). The last one was incubated with 1% FBS solution to block nonspecific interactions for 30 min at RT with a DyLight 488-conjugated TRA-1-60 antibody (Thermo Fisher Scientific) for extracellular staining. This was followed by a wash with DPBS. Histograms were evaluated using MA900 (Sony Biotechnology, Japan).

### Karyotyping

Karyotype analysis using G-banding was conducted at the 7th passage in the Research Centre for Medical Genetics, Moscow. 15 metaphases spreads was analysed: 13 metaphases consisted of 46 chromosomes, while 2 metaphases displayed abnormal count of 45 chromosomes. In this 2 cases, chromosomes 6 and 11 were absent, respectively. This discrepancy likely appeared during the sample preparation. The X and Y chromosomes were present in all metaphases.Thus, the percentage of normal metaphases was 87%.

### Trilineage differentiation

To evaluate functional pluripotency of ERCi003-A iPSC line, spontaneous differentiation into three germ layers was carried out with the formation of embryonic bodies (EBs). Briefly, at the confluence of 80% at the 13th passage, cells were washed with DPBS (Gibco) and treated with Versene solution (Paneco) for 6 min in 37ºC. Then, cells were gently detached by plastic scrapper (SPL, Korea) and replated to ultra-low attachment 6-well plate (Corning). Formation of EBs in suspension using KO-DMEM medium (DMEM/F12 (Gibco) containing 15% KnockOut™ Serum Replacement (Thermo Fisher Scientific) were observed during 7 days and then EBs were transferred on 0.1% gelatin coated plates (Sarstedt, Germany). Spontaneous differentiation was performed using DMEM/F12 medium with 10% FBS changing medium every two days. After 7 days the presence of three germ layer markers was evaluated by immunostaining antibodies at the concentrations listed in Table 1.

### Mutation analysis

The genomic DNA from hiPSCs was extracted with ExtractDNA Blood Kit (Evrogen, Russia). The region of interest in the *INS* gene was amplified via PCR using primers presented in Table 1. Amplification was performed in a 15 µL reaction volume and consisting of 8.7 µL nuclease free water (Evrogen), 0.1 µL SynTaq DNA Polymerase (Syngen, Russia), 1.5 µL buffer, 1.2 µL MgCl_2_, 0.5 µL forward primer, 0.5 µL reverse primer, 1.5 µL dNTP mix, 1 µL template DNA. PCR was performed using a MiniAmp Plus Thermal Cycler AB (Thermo Fisher Scientific). The thermal cycle was programmed for 5 min at 95 °C for initial denaturation, followed by 35 cycles of 15 s at 95°C for denaturation, 20 s at 60°C for annealing, 40 s at 72 °C for extension, and 3 min at 72°C for the final extension. PCR products were examined by electrophoresis at 200 V for 30 min in a 1% (w/v) agarose gel in 1 x TAE buffer (Evrogen) with ethidium bromide. The marker used a 1 kb DNA ladder (Thermo Fisher Scientific). The electrophoresis gel was examined in UV light. In order to confirm the presence of mutation, the PCR products were analyzed by the Sanger method using 3500 Genetic Analyzer (Thermo Fisher Scientific).

### Short tandem repeat (STR) profiling

STR analysis was conducted on ERCi003-A iPSC line at passage 10 and parental HDFs. The amplification and evaluation of 20 autosomal STR loci (AMEL, D3S1358, TH01, D12S391, D1S1656, D10S1248, D22S1045, D2S441, D7S820, D13S317, FGA, TPOX, D18S51, D16S539, D8S1179, CSF1PO, D5S818, vWA, D21S11, SE33) was performed with COrDIS Plus STR Amplification Kit (Gordiz, Russia) (data are available on request from the authors).

### Mycoplasma testing

Absence of mycoplasma was assessed by PCR via MycoReport Kit (Evrogen, Russia) at the 5th and 12th passages, according to the manufacturer instruction.

## OBTAINING AND CHARACTERIZATIION OF THE CELL LINE

The passport and the full characteristics of the iPSC ERCi003-A cell line are presented in the Table 2 and 3, respectively.

**Table 2.** Passport of the human iPSC cell line ERCi003-A.

| Characteristics | Description |
| --- | --- |
| Unique stem cell line identifier | ERCi003-A |
| Institution | Endocrinology Research Centre |
| Approval of the Ethics Committee | Ethics Committee of Endocrinology Research Centre, Moscow, Russia, Protocol no. 13 of July 12, 2023 |
| Cell type | iPSC |
| Species of organism | Human |
| Additional origin info | Age: 53<br>Sex: male<br>Ethnicity if known: Russian |
| Initial cell type | Dermal fibroblasts |
| Date of biomaterial collection | September 15, 2023 |
| Reprogramming method | Non-integrating, self-replicating RNA-based reprogramming vector |
| Reprogramming factors | OCT4, KLF4, SOX2, GLIS1, c-MYC |
| Clonality | Clonal |
| Genetic modification | No |
| Type of genetic modification | No |
| Disease | Maturity-onset diabetes of the young type 10 |
| Gene/locus | *INS* gene located on chromosome 11p15.5, Mutation: c.93C>G, p.C31W |
| Morphology | Monolayer colonies similar to human pluripotent cells |
| Pluripotency | Confirmed in the test for the formation of embryoid bodies |
| Karyotype | 46,XY |
| Contamination check | Bacteria, fungi and mycoplasma were not detected |
| Area of application | *In vitro* MODY10 model |
| Cultivation method | Feeder free (Matrigel, Corning) |
| Culture medium | mTeSR™1 (StemCell Technologies) |
| Temperature, °C | 37 |
| Concentration CO<sub>2</sub>, % | 5 |
| Concentration O<sub>2</sub>, % | 20 |
| **Passaging method** | Non-enzymatic, Versene solution (Paneco) |
| **Multiplicity of dilution during passaging** | 1:15 – 1:20 |
| **Cryopreservation** | CryoStore (StemCell Technologies) |
| **Storage conditions** | Liquid nitrogen |
| **Account in the registry** | https://hpscreg.eu/cell-line/ERCi003-A |
| **Date archived/stock date** | July 06, 2024 |

**Table 3.** Characterization and validation.

| Classification | Test | Result | Data |
| --- | --- | --- | --- |
| Morphology | Photography Bright field | Normal | Fig.1 panel a |
| Phenotype | Qualitative analysis:<br>Immunocytochemistry | Positive for pluripotency markers: OCT4, SOX2, TRA-1-60. | Fig.1 panel d |
|  | Quantitative analysis:<br>Flow cytometry | Expression of pluripotency marker TRA-1-60. Percentage of positive cells: 97.7% (n = 50.000) | Fig.1 panel b |
| Genotype | Karyotype (G-banding) and resolution | 46,XY, Resolution 450-500 | Fig.1 panel c |
| Identity | STR analysis | 20 loci analyzed, all matched | Data are available on request from the authors |
| Mutation analysis | Sequencing | Heterozygous [c.93C>G, p.C31W] in the *INS* gene | Fig.1 panel f |
| Microbiology and virology | Mycoplasma | Mycoplasma testing by PCR. Negative | Fig.1 panel g |
| Differentiation potential | Embryoid body formation | Positive staining for germ layer markers: brachyury and desmin (mesoderm), NEUROD1 and TUBB3 (ectoderm), FOXA2 and CXCR4 (endoderm) | Fig.1 panel e |

Upon signature of an informed consent, HDFs were derived from skin biopsy of a 53-year-old male carrying the mutation. HDFs were reprogrammed by transfection with non-integrating, self-replicating RNA vector encoding five reprogramming factors OCT4, KLF4, SOX2, GLIS1, c-MYC and a puromycin-resistant cassette in a single construct. To determine optimal concentration of puromycin dihydrochloride in order to select successfully transfected fibroblasts antibiotic titration firstly was performed. Optimal concentration of puromycin was 0.36 µg/ml. After 21 days, post-transfection observed colonies with embryonic stem cell (ESC)-like morphology were picked and expanded within the following days. Obtained ERCi003-A cell line have shown typical ESC-morphology – cells had a high nucleus to cytoplasm ratio and tightly packed structure (Fig. 1a). The pluripotency of ERCi003-A cells was confirmed by immunofluorescence staining with antibodies against OCT4, SOX2 and TRA-1-60 (Fig. 1b and d). The cytogenetic analysis of the obtained iPSC line showed a normal diploid (46,XY) profile (Fig. 1c). We evaluated the functional pluripotency of the obtained cells by their ability to form embryoid bodies and differentiate into the three germ layers. This was demonstrated by the positive expression of mesodermal markers – T-Box Transcription Factor T (brachyury) and desmin, ectodermal markers – tubulin beta-3 chain (TUBB3) and neurogenic differentiation factor 1 (NEUROD1), and endodermal markers – forkhead box protein A2 (FOXA2) and C-X-C chemokine receptor type 4 (CXCR4), as observed through immunofluorescence microscopy (Fig. 1e). Sanger sequencing of ERCi003-A cells confirmed the presence of pathogenic heterozygous mutation in the INS gene (Fig. 1f). In addition, short tandem repeat (STR) analysis was performed to test the allele match between ERCi003-A and fibroblasts. ERCi003-A cell line was free of mycoplasma contamination (Fig. 1g) (See Table 3).

**Fig 1.**
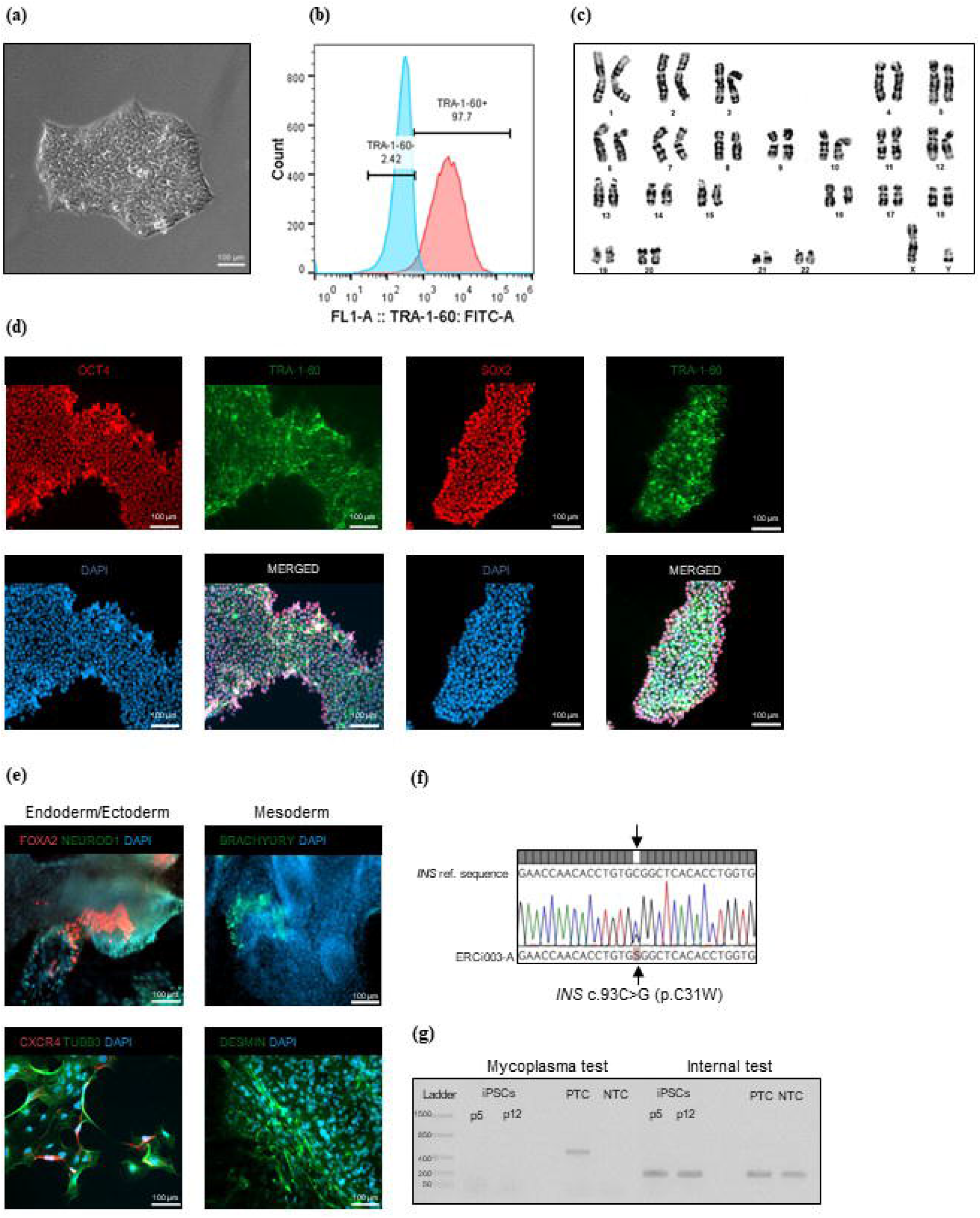
Characterization of the ERCi003-A human iPSC line. (a) cell morphology; (b) TRA-1-60 marker analysis by flow cytometry at the 13th passage; (c) karyotype analysis using G-banding at the 7th passage showed normal karyotype (46,XY); (d) immunofluorescent staining for pluripotency markers: OCT4 (red), TRA-1-60 (green), SOX2 (red) the 14th passage; (e) immunofluorescent staining for markers of three germ layers at the 13th passage: endoderm (FOXA2 (red), CXCR4 (red)), ectoderm (NEUROD1 (green), TUBB3 (green)), mesoderm (brachyury (green), desmin (green)); (f) Sanger sequencing result: *INS* gene polymorphism in iPSC line; (g) results of PCR analysis: no contamination of the line with mycoplasmas was detected at the 5th and 12th passages. All scale bars are 100 microns.

## FUNDING

This research was funded by the Ministry of Science and Higher Education of the Russian Federation (Grant Number: 075-15-2022-310).

## COMPLIANCE WITH ETHICAL STANDARDS

### Conflict of interest

The authors declare that they have no known competing financial interests or personal relationships that could have appeared to influence the work reported in this paper.

### Statement of compliance with standards of research involving humans as subjects

The study was approved by the Ethics Committee of Endocrinology Research Centre, Moscow, Russia, Protocol no. 13 of July 12, 2023. The patient was provided with all information about this study and signed an informed consent and an information sheet with his own hand.

## AUTHORS CONTRIBUTION

All authors have made substantial contributions to the conception and design of the study, or acquisition of data, or analysis and interpretation of data. In detail Asya Bastrich designed, performed experiments of obtaining the cell line and wrote the manuscript; Daniil Antonov performed experiment on characterization of cell line and analysed obtained data; Aleksandra Podzhilkova wrote the manuscript and analysed obtained data; Darya A. Petrova and Svetlana V. Pylina provided experimental set-up for the characterization assays; Dmitriy N. Laptev, Elena A. Sechko and Sergey N. Kuznetsov carried out medical support of the patient and the provision of dermal fibroblasts; Ekaterina A. Vetchinkina performed mutation analysis; Natalia G. Mokrysheva led and supported the project. All the authors revised the manuscript and approved the final version to be submitted.

## ACKNOWLEDGEMENT

Collecting of biological material of a patient with a pathological variant of *INS*: c.93C>G (p.C31W), isolation of dermal fibroblasts and genotyping, reprogramming of fibroblasts and characterization of iPSCs, immunofluorescence imaging was performed using resources of the Endocrinology Research Centre. Karyotype analysis was provided by the Research Centre for Medical Genetics. STR analysis was provided by the Gordiz Company.

## Notes

### Competing Interest Statement

The authors have declared no competing interest.

